# The chemical landscape of the human ribosome at 1.67 Å resolution

**DOI:** 10.1101/2023.02.28.530191

**Authors:** Alexandre Faille, Kyle C. Dent, Simone Pellegrino, Pekka Jaako, Alan J Warren

## Abstract

The ability of ribosomes to translate the genetic code into protein requires a finely tuned ion and solvent ecosystem. However, the lack of high-resolution structures has precluded accurate positioning of all the functional elements of the ribosome and limited our understanding of the specific role of ribosomal RNA chemical modifications in modulating ribosome function in health and disease. Here, using a new sample preparation methodology based on functionalised pristine graphene-coated grids, we solve the cryo-EM structure of the human large ribosomal subunit to a resolution of 1.67 Å. The accurate assignment of water molecules, magnesium and potassium ions in our model highlights the fundamental biological role of ribosomal RNA methylation in harnessing unconventional carbon-oxygen hydrogen bonds to establish chemical interactions with the environment and fine-tune the functional interplay with tRNA. In addition, the structures of three translational inhibitors bound to the human large ribosomal subunit at better than 2 Å resolution provide mechanistic insights into how three key druggable pockets of the ribosome are targeted and illustrate the potential of this methodology to accelerate high-throughput structure-based design of anti-cancer therapeutics.

## MAIN

Human ribosomes are complex nanomachines that translate the genetic code into cellular protein. They comprise 80 ribosomal proteins and four ribosomal RNAs (rRNAs) that carry a number of post-transcriptional modifications, most commonly 2’-OH ribose methylation or the isomerisation of uridine to pseudouridine^1^. rRNA modifications are an important source of ribosome heterogeneity that may influence ribosome function in response to environmental or developmental cues or in disease states^2^. However, the lack of high-resolution human ribosome structures determined using either X-ray crystallography or electron cryo-microscopy (cryo-EM) approaches, with accurately modelled potassium (K^+^), magnesium (Mg^2+^) ions and water molecules, is a significant barrier to more fully understand ribosome function and the fundamental structural role of ribosome modifications. The recognition that cancer cells are addicted to protein synthesis^3^ has further stimulated interest in generating high-resolution ribosome structures to accelerate the structure-based design of ribosome inhibitors as anti-cancer therapeutics.

The resolution of ribosome cryo-EM reconstructions is generally limited by thick ice and the interaction of ribosomes with the air-water interface during sample vitrification, which results in biased orientation distribution or particle damage^4,5^. Here, inspired by recent innovations in cryo-EM sample preparation and grid functionalisation^6,7^, we have overcome these limitations to determine the structure of the human 60S ribosomal subunit at a resolution of 1.67 Å. Our map allowed us to directly visualise thousands of water molecules, distinguish Mg^2+^ and K^+^ ions which are critical for the structural integrity and function of the ribosome, identify bound polyamine and unambiguously assign 117 rRNA modifications. The structure reveals the fundamental biological role of rRNA methylation in promoting carbon-oxygen hydrogen interactions (CH···O) to stabilise key functional areas of the ribosome. By revealing how translational inhibitors (blasticidin S, cycloheximide, and homoharringtonine) target the three key druggable pockets of the human 60S subunit, we highlight the potential for our rapid, simple, and inexpensive method to accelerate structure-based drug design on the ribosome.

### Improving human 60S particle orientation distribution and grid preparation

To improve the resolution of our human 60S subunit cryo-EM maps, we designed a methodology that improves particle orientation distribution and reduces both interaction with the air-water interface and ice thickness. We hypothesised that binding of the anti-association factor eukaryotic initiation factor 6 (eIF6) to purified human 80S-derived 60S ribosomal subunits might reduce over-represented views of the ribosome and increase the sampling of particle orientational space (**Fig. 1a**). Indeed, this strategy improved the particle orientation distribution, thereby minimising the number of micrographs required to obtain a high quality dataset (**Fig. 1b**). As an alternative to glow-discharge, we next developed an approach to improve the adsorption of biological specimens onto cryo-EM grids. We homogeneously functionalised pristine graphene-coated grids (or 2 nm carbon-coated grids) using the pyrene derivative 1-pyrenemethylamine (PMA) (**Fig. 1a**), rationalising that the positively charged amine function of PMA (predicted pKa 8.3 by QSAR modelling in OPERA^8^) would enhance binding of the negatively charged rRNA of the 60S subunit to the surface of the grid, thus reducing contact with the deleterious air-water interface. Using electron cryo-tomography, we determined the ice thickness of the grid sample to be 28 nm (**Fig. 1c**), equivalent to the largest dimension of the 60S subunit. By analysing refined defocus values on a per-particle basis within the micrographs, we estimated that the majority of 60S subunits lay within a 20-60 nm vertical ‘z’ range (median 39 nm across 9936 micrographs) (**Fig. 1d**), suggesting that optimal thin ice is present throughout the PMA-treated grids. By contrast, this distance extended to 30-90 nm (median 62 nm across 2260 micrographs) for the same sample when prepared using standard glow-discharged 2 nm carbon-coated grids. Furthermore, the homogeneity of the PMA-treated grids (**Supp. Fig. 1a**) allowed for automated acquisition of high contrast images from isolated particles (**Supp. Fig. 1b**), 97% of which were suitable for computing the final 3D reconstruction. Consequently, compared to glow-discharged grids, Rosenthal-Henderson ‘B-factor’ plots^9^ for datasets generated from PMA-treated grids were significantly improved, resulting in higher resolution reconstructions and enhanced throughput (**Fig. 1e-f and Supp. Fig. 1c-d**). Using this optimised methodology, we resolved a cryo-EM map of the human 60S ribosomal subunit to an overall resolution of 1.67 Å, generating the most accurate model of the human 60S ribosomal subunit to date (**Supp. Table 1** and **Supp. Fig. 2a, b**).

**Figure 1:**
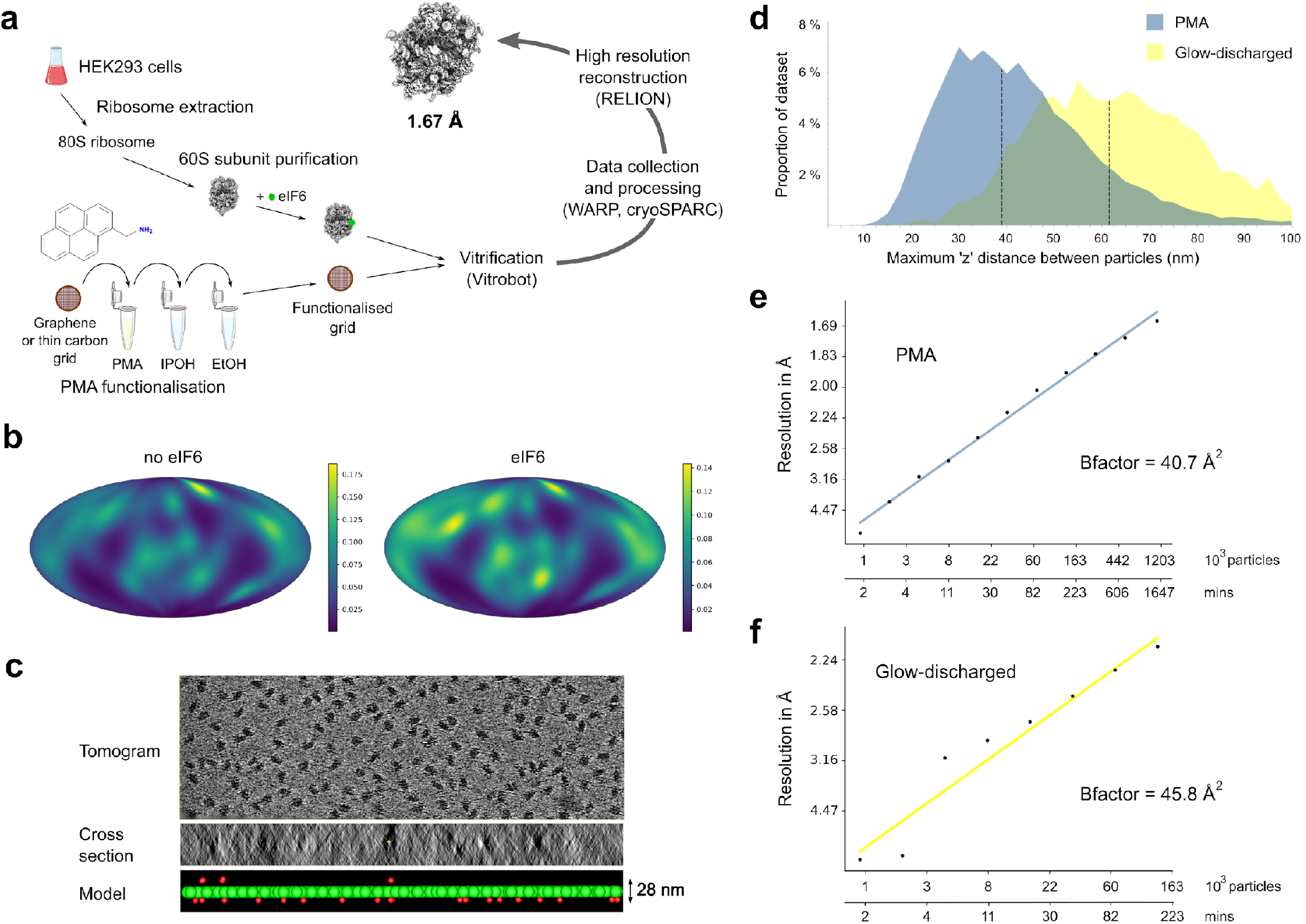
Strategy to improve cryo-EM data quality for the human 60S ribosomal subunit. **a**. Schematic of PMA-based sample preparation method and data processing to achieve high resolution. PMA, 1-pyrenemethylamine. IPOH, isopropanol. EtOH, ethanol. **b**. Orientation distribution of particles from the PMA dataset without (left) and with (right) eIF6 bound. Scale bar shows kernel density of particles across the orientation sphere. **c**. Representative cryo-tomogram from the PMA-treated grid and its cross-section used to measure ice thickness. Ribosome particles, green spheres; ice contaminants, red spheres. Ice thickness (distance between the carbon layer on the grid to the air-water interface) is equivalent to the largest dimension of the 60S subunit. **d**. Distribution of all micrographs from the indicated datasets as a function of the range of refined defocus values. PMA (blue), 60S-homoharringtonine complex dataset generated using PMA-based method; glow discharged (yellow), apo-60S dataset generated using glow-discharged grids. **e-f**. Rosenthal-Henderson ‘B-factor’ plots for PMA (**e**) and glow discharged (**f**) datasets as a function of both the number of particles and time of data collection, recorded at a rate of 350 movies / h (See Methods).

### Direct assignment of solvent, including magnesium and potassium ions

Optimal protein synthesis depends on the presence of water molecules, polyamines, divalent Mg^2+^ ions and monovalent cations such as K^+10^. However, direct assignment of these components within the human ribosome has remained elusive due to the limited resolution of existing structures. The assignment of K^+^ ions has depended on measuring the anomalous signal for K^+^ at the K-edge, derived from long-wavelength X-ray diffraction data collected from bacterial ribosome crystals^11^ but such data is not available for eukaryotic ribosomes. In addition, reliance on anomalous signal without thoroughly evaluating the stereochemistry of bound ions, which is difficult in the absence of any signal for water molecules, may lead to inaccurate ion assignment^12^. Our high quality cryo-EM maps allowed us to directly assign 11962 water molecules, 230 Mg^2+^ ions and 155 K^+^ ions (**Fig. 2**, central panel). The systematic octahedral coordination of Mg^2+^ ions at an idiosyncratic distance of 1.8-2.2 Å^13^, allowed us to assign them unambiguously (**Supp. Fig. 2c**). We assigned K^+^ ions using strict stereochemical criteria (see **Methods**) and the high signal peak visible in the map (**Supp. Fig. 2d**). Encouraged by the observation of a discrete signal for hydrogen atoms in Fo-Fc difference maps (**Supp. Fig. 3a**)^14–16^, we used these to validate the assignment of Mg^2+^ and K^+^ ions (and rRNA modifications) (**Supp. Fig. 3b-d**) to further improve the quality of the atomic model for the 60S ribosomal subunit.

**Figure 2:**
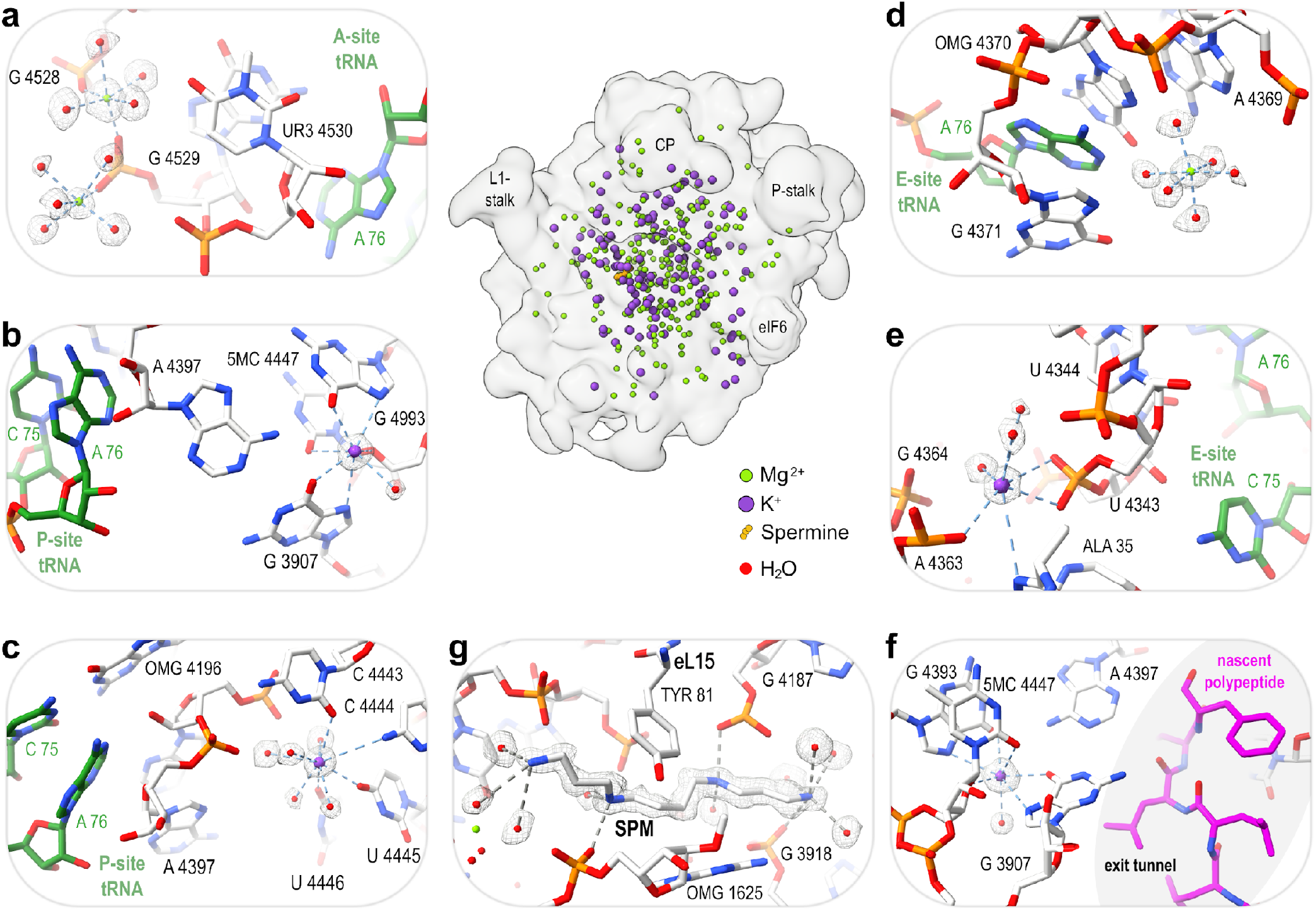
Direct observation of solvent molecules in functional sites of the human 60S ribosomal subunit. **Central panel**. Distribution of Mg^2+^ ions, K^+^ ions and spermine in the 60S subunit. CP, central protuberance. **a-f**. Mg^2+^ and K^+^ ions stabilise functional rRNA residues in tRNA binding sites. Previously solved structures of tRNAs (green) were superimposed on the human 60S ribosomal subunit structure in the A- (**a**), P- (**b, c**) and E-sites (**d, e**) and a nascent polypeptide (magenta) superimposed in the exit tunnel (**f**) using PDB IDs 6OLF, 6SGC and 5LZV. **g**. Spermine (SPM) interacts with ribosomal protein eL15, rRNA and waters in a binding pocket ∼20 Å from the polypeptide exit tunnel. Ion coordination (blue dashed lines); hydrogen bonds (grey dashed lines); OMG, 2’-O-methylguanosine; 5MC, 5-methylcytosine; CP, central protuberance.

Similar to Mg^2+^ ions, K^+^ ions bind throughout the 60S subunit and play an essential role in maintaining structural and functional integrity. Superimposing human (A-site, PDB 6OLF) and rabbit (P-site, PDB 6SGC; E-site, 5LZV) tRNA bound complexes reveals how K^+^ and Mg^2+^ ions create a network of interactions with the surrounding rRNA and water molecules to fine-tune the position of rRNA residues responsible for tRNA binding (**Fig. 2a-e**). Similarly, K^+^ and Mg^2+^ ions also stabilise rRNA residues close to the polypeptide exit tunnel, within reach of the nascent polypeptide (**Fig. 2f**).

Despite the importance of polyamines for the structural stability of ribosomes and for translation *in vivo*^17^, to date they have only been modelled with low confidence and accuracy in eukaryotic ribosomes^18,19^. We unequivocally identified a spermine molecule bound within the core of the human 60S subunit, in a cleft formed by rRNA helices 32, 33, 74, and 75 and the ribosomal protein eL15 (**Fig. 2g**). Together with the phosphate group of residue G 3918, multiple water molecules stabilise the extremities of the spermine molecule, while residues G 1625 and G 4187 make hydrogen bonds with its inner amine functions. The side chain of residue Tyr 81 of ribosomal protein eL15 and the nucleobase of G 1625 further anchor the spermine molecule through stacking interactions. As no exogenous spermine was added during sample preparation, the identified spermine molecule is likely an integral structural component of the human ribosome that helps maintain local stability by linking disparate structural elements.

### Functional role of ribosomal RNA modifications

The accurate identification of post-transcriptional ribosome modifications is critical for a complete understanding of their role in ribosome biogenesis, translation and human disease^20^. We directly visualised 117 of the 137 rRNA modifications identified by quantitative mass spectrometry^21^ in the human 60S subunit (**Fig. 3, Supp. Table 2, Supp. Fig. 4a-b**), leveraging Fo-Fc difference maps to confirm modifications of both the rRNA (**Supp. Fig. 3d**) and ribosomal proteins^22^ (**Supp. Fig. 3e**). Of the remaining 20 rRNA modifications identified by mass spectrometry^21^, four could not be visualised despite their location in well-defined areas of the cryo-EM map, consistent with the suggested heterogeneity of some non-essential modifications^23,24^. The remaining 16 modifications were not visualised due to low local resolution of the map. Proposed human-specific modifications based on cryo-EM^25^ but not confirmed by mass spectometry^21^, were not identified in our map (**Supp. Fig. 4c**).

**Figure 3:**
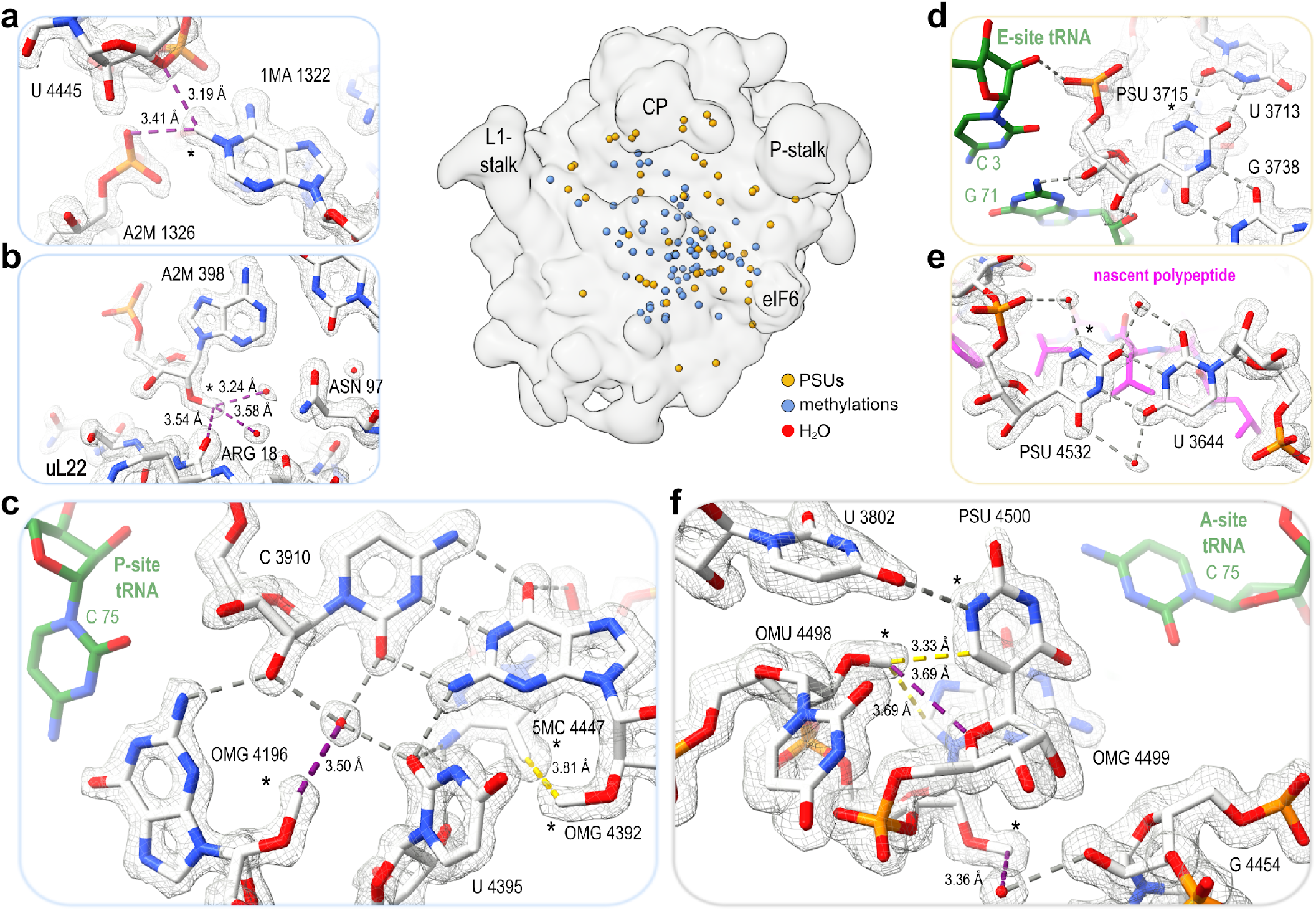
Role of rRNA modifications in the human 60S ribosomal subunit. **Central panel**. Distribution of rRNA modifications within the 60S ribosomal subunit. **a-c**. Structural role of rRNA methylation. Methyl groups form CH···O interactions (purple dashed lines) with phosphate (**a**), water, and protein (**b**). **c**. Functional role of rRNA methylation in the P-site. CH···O hydrogen bonding interaction with a water molecule helps orientate OMG 4196 to interact with the P-site tRNA (green), superimposed from PDB ID 6SGC. Hydrophobic interaction between the 5’-methyl group of 5MC 4447 and the 2’-O-methyl group of OMG 4392 is shown as a yellow dashed line. **d-e**. Functional role of nucleoside pseudouridylation in the 60S ribosomal subunit. PSU 3715 helps maintain the position of the E-site tRNA acceptor stem (green), superimposed from PDB ID 5LZV (**d**); PSU 4532 helps to delineate the proximal part of the polypeptide exit tunnel (nascent polypeptide, magenta), superimposed from PDB ID 6OLF (**e**). **f**. Methylated residues OMU 4498, OMG 4499, and PSU 4500 combine hydrophobic and hydrogen bonding (including CH···O bonds) interactions to stabilise the A-site tRNA binding site, tRNA structure superimposed from PDB ID 6OLF. OMG, 2’-O-methylguanosine. 5MC, 5-methylcytosine. PSU, pseudouridine. *, rRNA modifications.

We next sought to use our high quality maps with accurately assigned solvent molecules to better understand the biological role of rRNA methylation. For 60 of the 66 methylated rRNA residues observed (**Supp. Table 2**), the extra methyl group establishes unconventional CH···O hydrogen bonding interactions^26–32^, mostly with water molecules but also with phosphate or ribose moieties from neighbouring rRNA residues, or with protein partners (**Fig. 3a-f**). While most methylations appear to have a general structural role, others likely modulate ribosome function. For example, the direct interaction of residue OMG (2’-O-methylguanosine) 4196 with the CCA end of the P-site tRNA (residue C 75) is stabilised by the CH···O interaction formed between the 2’-O-methyl group of residue OMG 4196 and a highly stable water molecule (B-factor of 5 Å^2^) that connects it to a wider network of rRNA interactions (**Fig. 3c**). Also within this network, there is a direct hydrophobic interaction between the methyl groups of residues 5MC (5-methylcytosine) 4447 and OMG 4392 that further stabilises this area. Thus, rRNA methylation exploits both unconventional CH···O and hydrophobic interactions to extend the reach of rRNA residues with the chemical environment of the ribosome.

We unambiguously identified 52 of the 63 pseudouridines (PSU) indicated by mass spectrometry^21^ by directly observing the hydrogen bonds formed by the N1 atom (**Supp. Fig. 4b**). Similar to rRNA methylations, the hydrogen bonding potential of pseudouridine compared to uridine is exploited to position and stabilise rRNA residues with key functional roles. For example, residue PSU 3715 interacts with residue U 3713 through its N1 atom and stabilises the E-site tRNA through interactions with residues C 3 and G 71 (**Fig. 3d**). Pseudouridine N1 atoms can also form hydrogen bonds with water molecules. For example, a water molecule bridges the phosphate group and base of PSU 4532, thereby helping to delineate the polypeptide exit tunnel (**Fig. 3e**). rRNA methylation and pseudouridylation works in a synergistic manner, whereby methylated residues OMU (2’-O-methyluridine) 4498 and OMG 4499 cooperate with PSU 4500 to form the tRNA A-site binding pocket through a combination of hydrophobic interactions and hydrogen bonds, including CH···O bonds (**Fig. 3f**).

### Binding mechanism of inhibitors targeting the human 60S ribosomal subunit

We next tested the applicability of our methodology to accelerate structure-guided drug design on the 60S ribosomal subunit. Using PMA-treated grids, we determined cryo-EM structures of the translational inhibitors blasticidin S, cycloheximide and homoharringtonine bound to the human 60S subunit at resolutions of 1.86 Å, 1.90 Å and 1.67 Å, respectively (**Fig. 4, Supp. Fig. 5**). At an acquisition rate of around 350 micrographs / h, the high signal-to-noise ratio observed for our samples allowed us to determine high resolution maps using less than 4 h of microscope time **(Supp. Fig. 6)**.

**Figure 4:**
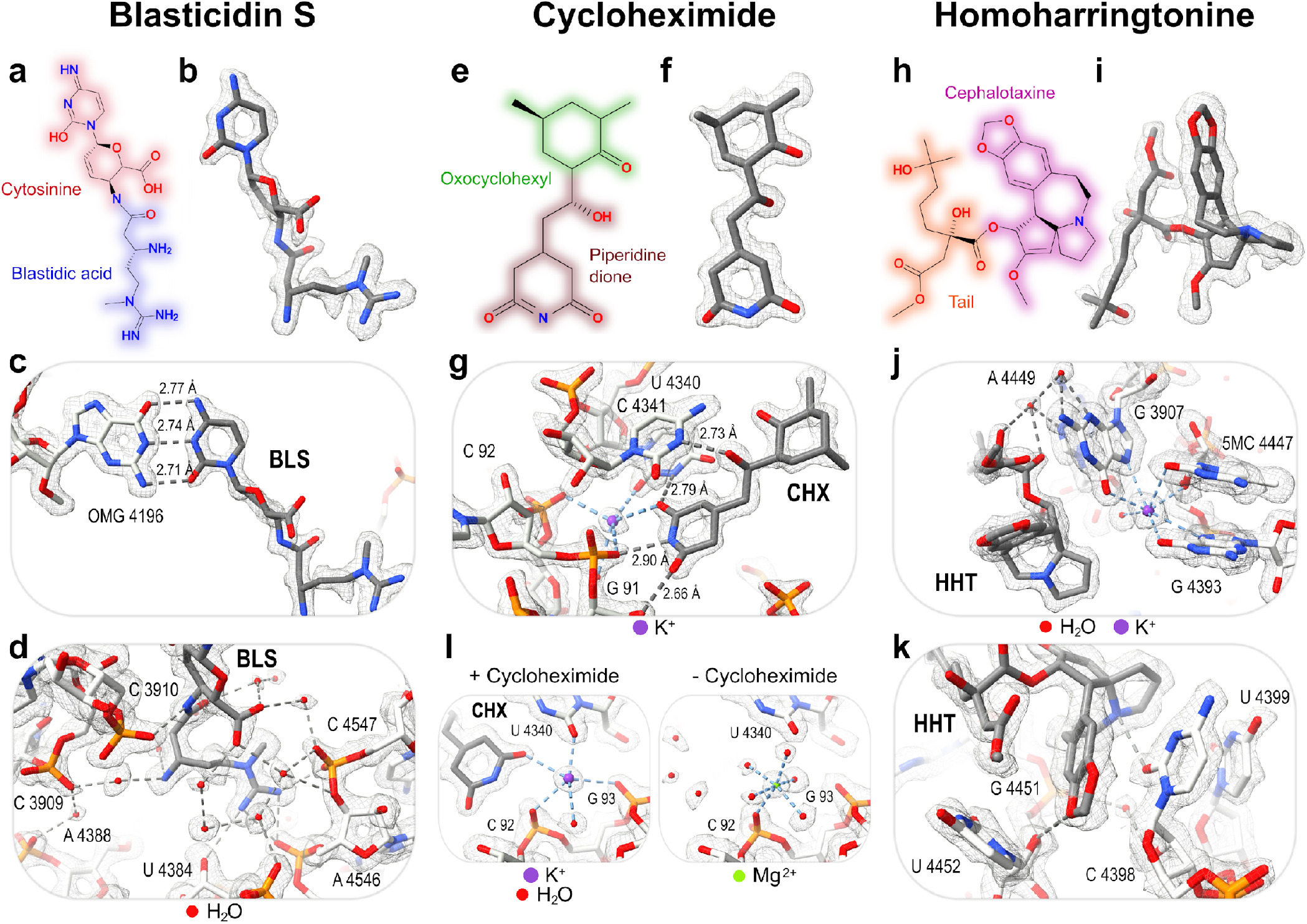
Mechanisms of inhibitor binding to the human 60S ribosomal subunit. **a-l**. 2D chemical structures fit into the cryo-EM map and interactions with the chemical environment for blasticidin S (**a-d**), cycloheximide (**e-g**) and homoharringtonine (**h-k**). **l**. Comparison of cycloheximide-bound and cycloheximide-free structures showing a K+ ion co-bound with cycloheximide in the E-site tRNA binding pocket. BLS, blasticidin S. CHX, cycloheximide. HHT, homoharringtonine.

Blasticidin S forms short (2.7-2.8 Å) hydrogen bonds with the 2’-O-methylated base G 4196 that mimic Watson-Crick base pairing (**Fig. 4a-c**), in contrast with recent suggestions^33^. Blasticidin S thus binds the ribosome at the same position as residue C 75 from the P-site tRNA, outcompeting it to block translation (compare with **Fig. 2c**). A network of water molecules interacts with blasticidin S to stabilise the cytosinine moiety (**Fig. 4d**) and pulls the primary amine of the blastidic acid moiety closer to base C 3909, forming a cleft into which the phosphate group of A 4388 enters. In turn, the tip of the blastidic acid moiety forms multiple electrostatic interactions with the phosphate groups of rRNA residues A 4546 and C 4547, and the ribose 2’-OH of U 4384.

Cycloheximide forms three short hydrogen bonds, one (2.73 Å) between its hydroxyl group and base C 4341 (atom N3) and two from its piperidine-dione moiety to bases G 91 (2’-O, 2.66 Å) and U 4340 (atom N3 atom, 2.79 Å) (**Fig. 4e-g**). The piperidine-dione moiety of cycloheximide interacts with a K^+^ ion stabilised by the phosphate backbone residues of bases G 91 and C 92 (**Fig. 4g**). This K^+^ ion plays a central role in the binding of cycloheximide to the ribosome but was previously assigned as a Mg ^2+^ ion^34,35^. Interestingly, a Mg^2+^ ion is found *in lieu* of K^+^ in the absence of cycloheximide (**Fig. 4l**).

Homoharringtonine binds within the A-site cleft, in a pocket delineated by residues G 3907, G 4393, and A 4449, which are stabilised through interactions with a K^+^ ion (**Fig. 4h-j**). A single well-defined water molecule connects residues A 4449 and G 3907 with the homoharringtonine tail through hydrogen bonding interactions (**Fig. 4j**). Together with the interaction with residue U 4452, this allows stacking between residue C 4398, the tail, and the cephalotaxine moiety of homoharringtonine (**Fig. 4k**).

The most accurate structures of these translational inhibitors to date are for complexes with the *S. cerevisiae* ribosome^35^. However, our maps reveal key differences in the position and conformation of all three inhibitors bound to the human 60S subunit compared with yeast. In the human 60S-inhibitor complex, the cytosinine group of blasticidin S aligns with residue OMG 4196, closely mimicking Watson-Crick G-C base pairing (**Supp. Fig. 7a**), while in yeast, the cytosinine group is misaligned with residue OMG 4196 as it is tilted 31 degrees towards residue G 4197 (human rRNA numbering). Compared to yeast, in the human 60S-cycloheximide complex, ribosomal protein eL42 (residues 52-59) is positioned closer to cycloheximide, likely due to electrostatic interactions between Arg57 (Phe in yeast) and the phosphate backbone of the rRNA (**Supp. Fig. 7b**). Finally, compared to its yeast counterpart, the conformation of the homoharringtonine tail bound to the human ribosome also differs (**Supp. Fig. 7c**). Taken together, our structures provide important new insights into the mechanisms of inhibitor binding to the ribosome and highlight the critical need to visualise human ribosome-inhibitor complexes to design new anti-cancer drugs with higher affinity and specificity.

## DISCUSSION

In this study, we report the development of a robust methodology that has allowed us to determine the structure of the human 60S ribosomal subunit to a resolution of 1.67 Å (map-to-model FSC: 1.68 Å), one of the highest resolution cryo-EM structures ever deposited in the Protein Data Bank and on par with the recently determined structure of the *E. coli* ribosome (half-maps FSC: 1.55 Å; map-to-model FSC: 1.73 Å; PDB code 8B0X). Our structure represents the most complete and accurate atomic model of the human 60S ribosomal subunit to date, including water molecules, K^+^ and Mg^2+^ ions, spermine and rRNA chemical modifications, with bound eIF6, providing a key reference for future studies of ribosome biogenesis, protein synthesis and human ribosomopathies^36^.

Our findings reveal the fundamental, conserved biological role of rRNA methylation in forming a widespread network of CH···O interactions that promote stability by increasing the range of potential interactions within the chemical environment of the ribosome. The majority (91%) of methylated rRNA residues are involved in CH···O interactions, not only with the waters, ribose or phosphate moieties from neighbouring rRNA residues, but also with the backbone residues of ribosomal proteins. Furthermore, the direct visualisation of rRNA modifications and the new insight into their functional role will help inform the biology of cancer predisposition disorders such as dyskeratosis congenita where the chemical nature or stoichiometry of such modifications is altered^2,37^. By accurately assigning the solvent shell of the ribosome, our structure highlights the functional role of K^+^ and Mg^2+^ ions in fine-tuning the position of rRNA residues both at key tRNA binding sites and within the peptide exit tunnel.

The structures of the translational inhibitors homoharringtonine, cycloheximide, and blasticidin S bound to the human 60S subunit reveal the detailed mechanism by which the three main druggable pockets of the ribosome, the tRNA binding sites, are targeted. The structures accurately position the ligands and provide precise distance measurements for the interactions within the ribosome, including ions and solvent, that together drive the affinity and specificity of inhibitor binding. While sub-2 Å resolution blasticidin S and cycloheximide datasets were obtained in less than 4 hours, reconstructions at 2.5 Å resolution required less than 15 min of data collection (**Fig. 1e, Supp. Fig. 6a**) while still generating high quality maps (**Supp. Fig. 8**). The high signal peak in Fo-Fc difference maps produced by unmodelled molecules in such maps allows the rapid detection and identification of bound compounds. Homogeneous ice thickness and sample dispersion across the grid (**Fig. 1d, Supp. Fig. 1a-b** and **Supp. Fig. 6b**) permit automated square picking algorithms to be used with minimal user intervention, further increasing throughput. Our strategy is inexpensive, reproducible, and relies on standard cryo-EM equipment. Due to the gain in achievable resolution and high throughput of the approach, our methodology represents a step change for the rational structure-based design of new and improved drugs targeting the human ribosome.

## MATERIALS AND METHODS

### Human 60S ribosomal subunit purification

Human 60S ribosomal subunits were prepared as described, with some modifications^38^. Expi293F^™^ human cells (ThermoFisher Scientific) were grown in Expi293^™^ medium (Gibco) at 37 °C and 8 % CO_2_. Prior to harvesting cells in exponential growth, NaCl (final concentration 100 mM) was added to the medium for 20 min. Cells were centrifuged for 10 min at 600 x g, resuspended in phosphate-buffered saline (PBS) and centrifuged for 10 min at 600 x g. Cells were snap frozen in liquid nitrogen and stored at −80 °C. The cell pellet was thawed on ice, resuspended in lysis Buffer S (30 mM Hepes pH 7.5, 7.5 mM MgCl_2_, 50 mM KCl, 0.5 mM EDTA, 220 mM sucrose, supplemented with 0.5 mM NaF, 0.1 mM Na_3_VO_4_, one complete protease inhibitor tablet (Roche), 1 mg/mL final concentration of Na-heparin, 1 U/µL Rnasin® Plus RNAse inhibitor (Promega), 2 mM DTT, 0.5% final concentration of IGEPAL CA-630 (Sigma)) and incubated at 4 °C for 30 min with continuous mixing. The sample was centrifuged sequentially at 10,000 x g for 5 min and 30,000 x g for 20 min. PEG 20,000 (final concentration 1.35 %) was added for 10 min at 4 °C and centrifuged at 17,500 x g for 13 min. The pellet was discarded and the KCl concentration adjusted to 125 mM. Following incubation with PEG 20,000 (final concentration 5%) for 15 min at 4 °C, the solution was centrifuged at 17,500 x g for 11 min. The supernatant was discarded and the ribosome-containing pellet resuspended in buffer E (30 mM Hepes pH 7.5, 6 mM MgCl_2_, 125 mM KCl, 2 mM DTT) supplemented with 1 mg/ml Na-heparin. The solution was centrifuged over a 10-30 % sucrose gradient (30 mM Hepes pH 7.5, 7.5 mM MgCl_2_, 125 mM KCl, 0.5 mM EDTA) at 42,800 x g for 15 h using a SW28 rotor (Beckman Centrifuge). Gradients were fractionated using an AKTA system and the 80S ribosome fractions pooled. PEG 20,000 (5.25 % final concentration) was added to the pool for 10 min at 4 °C and the sample centrifuged at 17,500 x g for 10 min. The pellet was resuspended in buffer W (30 mM Hepes pH 7.5, 6 mM MgCl_2_, 100 KCl, 2 mM DTT) and centrifuged over a 10-40% sucrose gradient prepared in ribosome dissociating conditions (30 mM Hepes pH 7.5, 5 mM MgCl_2_, 550 mM KCl, 2 mM DTT) at 70,000 x g for 20.5 h using a SW28 rotor. 60S ribosomal subunit fractions were pooled and PEG 20,000 (6.23% final concentration) added. Ribosomes were then incubated for 15 min at 4 °C and precipitated at 17,500 x g for 10 min. The pellet was gently resuspended in buffer F (20 mM Hepes pH 7.5, 5 mM MgCl_2_, 100 mM KCl, 2 mM DTT), cleared at 15,000 x g for 5 min, aliquoted, and snap frozen in liquid nitrogen.

### eIF6 binding

eIF6-bound human 60S subunits were obtained by incubating purified 60S with exogenous eIF6 at 37 °C for 15 min followed by centrifugation over a 30% sucrose cushion prepared in buffer F and resuspension of the pelleted 60S subunits with buffer F. Exogenous eIF6 was obtained as described^39^. The amount of eIF6 added was optimised empirically to be in slight excess over 60S subunits. Briefly, mixes at various 60S:eIF6 ratios were centrifuged over a 30% sucrose cushion prepared in buffer F, the resulting pellet was resuspended in buffer F and subjected to immunoblotting using an anti-eIF6 antibody (GenTex GTX117971) to assess the amount of exogenous eIF6 required to saturate 60S subunits.

### Cryo-EM sample preparation

For PMA functionalisation, pristine graphene-coated R2/4 grids (Graphenea) were initially used for optimisation purposes but 2 nm formvar/amorphous carbon R2/2 grids (Quantifoil) were used for data collection. However, we recommend using graphene-coated grids which present great reproducibility, whereas 2 nm formvar/amorphous carbon ones can show significant batch-to-batch variability. Grids were processed at room temperature and handled individually using reverse tweezers, they were not glow discharged or otherwise plasma treated. Grids were soaked for 30 s in a freshly prepared 1 ml solution of 50mM 1-pyrenemethylamine (Sigma-Aldrich 401633) solubilised in DMSO. They were successively washed by soaking in 1 ml of isopropanol and 1 ml of ethanol, for 5 s each. Still suspended by tweezers, grids were allowed to air dry for 30 min. At the same time, eIF6-bound 60S subunits at a concentration of 120 nM were incubated with 100 µM inhibitors (final concentration) at 37 °C for 1 h. We added 1 µl of 10 mM DMSO-solubilised homoharringtonine (Sigma-Aldrich SML1091) to 100 µl of 60S subunits (homoharringtonine dataset), 1 µl of 2 mM water-solubilised blasticidin S (Sigma-Aldrich 15205) to 20 µl of 60S subunits (blasticidin S dataset) and 1 µl of 2 mM water-solubilised cycloheximide (Sigma-Aldrich 01810) to 20 µl of 60S subunits (cycloheximide dataset). Within the chamber of a Vitrobot Mark IV (Thermo Fisher Scientific) set at 100% humidity and 4 °C, 4 μl of 60S-inhibitor complex was deposited onto freshly prepared PMA-functionalised grids. Grids were blotted using the following parameters: blot time 1 s, blot force −7, wait time 30 s, no drain time. Blotted grids were vitrified in liquid ethane and stored in liquid nitrogen.

For the ‘glow-discharged’ (GD) dataset, eIF6-bound 60S subunits at 120 nM were deposited onto freshly glow-discharged grids (20 mA for 15 s using a PELCO easyGlow) and blotted using the same parameters as the PMA-functionalised grids.

### Ice-thickness determination by electron cryo-tomography

Specimens were prepared as for standard single-particle data-collection. Micrograph tilt-series were acquired using TOMO4/5 (Thermo Fisher Scientific) on a Talos Arctica (Department of Biochemistry, University of Cambridge) microscope operating at 200 keV using a Falcon 3 detector operating in linear mode. A nominal magnification of 45000 X (2.23 Å / pixel) was used to collect a tilt-series over a range of 45° using a step of 3° and total dose per tilt of 3.34 e^-^ / Å^2^ with a 0.25 s exposure time per tilt (a total dose per tilt-series of ∼93 e^-^ / A^2^).

Tilt-series were aligned using standard fiducial-less methods and down-sampled by a factor of four prior to reconstruction (8.92 Å / pixel) visualisation and modelling using the IMOD package v4.9 (University of Colorado, Boulder). Tomograms were calculated by back-projection and a SIRT-like filter applied (over 4 iterations) to improve ribosomal particle contrast and reveal the 2 nm formvar/carbon layer. The centres of ribosomal particles and of a limited number of contaminating ice clusters that decorated the surface of the ice-layer and opposing carbon layer were selected manually and modelled as spheres of approximately relevant sizes. Ribosomal spheres corresponded to a diameter of 28 pixels (∼25 nm). Ice clusters were modelled using a sphere size diameter of 14 pixels (∼12.5 nm diameter). An average ice thickness of 27.65 nm was estimated according to the distance between ice clusters on each side taking into consideration their own spherical radiuses.

### Cryo-EM single particle data collection and processing

Datasets were collected using Krios III at the Electron Bio-Imaging Centre (eBIC, Diamond Light Source, Harwell Science and Innovation Campus) operating at 300 keV. Electron micrographs were recorded using the ‘aberration free image-shift’ (AFIS) functions of EPU v2.5 (Thermo Fisher Scientific) using a Gatan Bio-Quantum K3 direct electron detector (Gatan, Pleasanton, USA). A nominal magnification of 105,000 X was used in EFTEM mode, using a slit-width of 20 eV and a pixel size of 0.829 Å / pixel (0.4145 Å / pixel in super-resolution mode). A total dose of 51 e ^-^ / A^2^ was achieved over a 2.2 second exposure (a dose rate of 15.97 e^-^ / px / s). A nominal defocus spread between −1.0 to −1.8 µm was specified with automatic focusing performed every 7.5 µm. The illumination diameter was 1.28 µm, allowing for 4 shots per ∼2.7 µm diameter specimen hole. Micrograph movies were recorded in TIFF format incorporating either 60 or 40 fractions, a dose per fraction of 0.85 e^-^ / Å^2^ /fraction or 1.275 e^-^ / Å^2^ / fraction, respectively.

Micrograph movies were corrected for beam-induced motion, and particles selected using the BoxNet neural network with low stringency using Warp^40^. Using an in-house script (‘Warp2Relion.py’), a RELION project was automatically built from the Warp session. Particles were extracted into a 512 pixel box using RELION and down sampled by a factor of four to improve contrast and signal-to-noise ratio. The particle stack was imported into CryoSPARC and initially reconstructed into 3-5 classes using the stochastic gradient descent or ‘ab initio’ method^41^. Stack cleaning was performed against these 3D references using ‘homogeneous refinement’ within CryoSPARC. Particle stack cleaning performed against 3D references, rather than a large number of 2D references, was shown to significantly improve particle representation for relatively unpopulated views of the complex. The CryoSPARC class of interest was then converted back into RELION STAR format using a custom implementation of Daniel Asarnow’s ‘pyem’ Python package (https://github.com/asarnow/pyem) that reintegrated the class back into the pre-existing RELION project.

From this point onwards, RELION^42,43^ was used for data processing. Movies were imported and motion corrected using MotionCorr2^44^ with dose weighting enabled and binning factor set to 2. Particles imported from CryoSPARC were extracted from the resulting micrographs in a 512 × 512 pixels box. 3D refinement was then performed using a mask of 320 Å diameter. The refined map was post-processed with a 15 Å low-pass mask generated using a binarisation threshold of 0.004, a binary extension of 2 pixels and a soft-edging of 5 pixels. Beamtilt, along with third and fourth order aberrations were first refined, followed by anisotropic magnification, and finally defocus, astigmatism, and B-factor values, on a per-particle basis. Bayesian polishing was then applied to the movies using recommended parameters. An identical cycle of 3D / CTF refinement was performed, followed by a final 3D refinement to generate the final map.

### Particle orientation distribution analysis

Particle orientation distributions were plotted with a script adapted from Naydenova *et al*.^45^ that employs a probability density function. 2D kernel density estimation (kde2d) was parameterised with a 100 × 100 bins grid and a bandwidth of 0.175°.

### Structure modelling and refinement

Structure modelling was performed in Coot^46^, building into a post-processed map generated in RELION (automatic sharpening of −40.7). As the global resolution of the dataset reaches Nyquist limit (0.83 A pixel size), the map was manually upsampled to 768 × 768 × 768 voxels with a corresponding pixel size of 0.5527 Å. PDB ID 6EK0 was used as a starting model, rRNA and protein chains, zinc and Mg2+ ions were visually inspected to correct for mismodelling or changes due to the absence of the small ribosomal subunit in our sample. rRNA modifications that could not be directly visualised were removed while others were added based on mass spectrometry data ^21^. Water molecules were automatically added in Coot, refined in REFMAC^47^, and further removed in Coot when not meeting the following criteria: minimum interaction distance 1.8 Å, max interaction distance 3.5 Å, maximum B-factor 70 Å^2^, minimum map rmsd level 5 e-/A^3^. Mg^2+^ and K^+^ ions that met established criteria^13^ were then added. For Mg^2+^ ions, these are a tight first coordination sphere with strict octahedral geometry and a coordination distance of 1.9-2.2 Å. For K^+^ ions, the criteria were a first coordination sphere comprising at least 5 coordinations and involving at least one oxygen atom (other than a water molecule) with a coordination distance of 2.7-3.2 Å and a signal peak in the map significantly higher than surrounding waters. We took a conservative approach such that whenever those criteria could not be fully met, we assumed that a highly coordinated and well-defined water was present and this was modelled accordingly. Finally, the resulting model was refined using a combination of PHENIX ^48^ and REFMAC and validated in REFMAC for 20 cycles using automatic weighting.

Methyl groups from rRNA modifications were considered to undergo CH···O hydrogen bonding if the C···O distance was <3.70 Å, the X–C···O angle was between 85° and 140°, and the elevation angle was <50°. The C···O–Y angle was required to be between 75° and 160°, according to published criteria^26,32^

Fo-Fc difference maps were calculated using Servalcat^16^ within CCP-EM^49^. When used to highlight hydrogen atom signals, default parameters were used (including 20 refinement cycles). When used to assist modelling, riding hydrogen atoms were kept through difference map calculation (keyword ‘make hout’). Whenever modifications were made to the final refined model, such as to highlight the presence of rRNA modifications or ions (*e*.*g*. Supp. Fig. 3b-e), 10 cycles of refinement were carried out in REFMAC prior to calculation of the difference map.

Molecular graphics and analysis were performed using UCSF ChimeraX^50^, developed by the Resource for Biocomputing, Visualization, and Informatics at the University of California, San Francisco, with support from National Institutes of Health R01-GM129325 and the Office of Cyber Infrastructure and Computational Biology, National Institute of Allergy and Infectious Diseases.

## AUTHOR CONTRIBUTION

A.F. contributed project design, experimental work, grid preparation, data processing, data analysis; K.C.D. contributed grid preparation, data collection, data pre-processing; S.P. and P.J. contributed experimental work; A.F. and A.J.W. wrote the manuscript with contributions from all authors.

## ACKNOWLEDGMENTS

We thank the group of Dr. V. Ramakrishnan for the gift of the Expi293 cells. This work was supported by Blood Cancer UK (21002), the UK Medical Research Council (MR/T012412/1), the Kay Kendall Leukaemia Fund, European Cooperation in Science and Technology (COST) Actions CA18233 “European Network for Innovative Diagnosis and treatment of Chronic Neutropenias, EuNet-INNOCHRON” and CA21154 “Translational control in Cancer European Network TRANSLACORE”, a Wellcome Trust strategic award to the Cambridge Institute for Medical Research (100140), a core support grant from the Wellcome Trust and MRC to the Wellcome Trust-Medical Research Council Cambridge Stem Cell Institute, the Cambridge National Institute for Health Research Biomedical Research Centre (BRC-1215-20014), the Connor Wright Project and the SDS Foundation, the Swedish Research Council (2014-06807, to PJ). We thank Dr. D. Y. Chirgadze for assistance with data collection at the Cryo-EM Facility, Department of Biochemistry, University of Cambridge, funded by the Wellcome Trust (206171/Z/17/Z; 202905/Z/16/Z), the Departments of Biochemistry and Chemistry, the Schools of Biological Sciences and Clinical Medicine and the University of Cambridge. We acknowledge Diamond Light Source for access and support of the cryo-EM facilities at the UK’s national Electron Bio-imaging Centre (eBIC) [under proposal EM BI22238], funded by the Wellcome Trust, MRC and BBRSC.

## DATA AVAILABILITY

Atomic coordinates of the human homoharringtonine-bound 60S subunit structure have been deposited in the Protein Data Bank with accession codes 8A3D, and the cryo-EM density maps have been deposited in the Electron Microscopy Data Bank with accession codes EMD-15113. Raw movies will be deposited to the Electron Microscopy Public Image Archive (https://www.ebi.ac.uk/pdbe/emdb/empiar/).

---

**Supplementary Table 1**: Data collection and refinement statistics

**Supplementary Table 2**: List of rRNA modifications and interaction partners in the human 60S ribosomal subunit The rRNA modifications identified in our cryo-EM map are compared to those previously reported using cryo-EM^25^ or mass spectrometry^21^. Compared to mass spectrometry results, modifications that could not be confirmed despite local high quality of the map are marked in red. Those that could not be confirmed due to low local map quality are denoted ND (not defined). The chemical nature of partners forming CH···O and hydrophobic interactions with the methyl group of rRNA methylations and those forming H-bonds with the N1 atom of pseudouridines is indicated.

## Supplementary Figures

