## Supplementary Table 1 for "The chemical landscape of the human ribosome at 1.67 Å resolution"

| | Homoharringtonine | Cycloheximide | Blasticidin S |
|---|---|---|---|
| **Data** | | | |
| Number of collected movies | 9973 | 1393 | 1401 |
| Initial particles images | 1147785 | 182359 | 180106 |
| Final particle images | 1088709 | 173946 | 159962 |
| Map resolution (Å) | 1.67 | 1.90 | 1.86 |
| Map sharpening b factor (Å$^2$) | 40.7 | 38.6 | 40.8 |
| **Model composition** | | | |
| Protein residues | 6578 | 6578 | 6578 |
| RNA residues | 3571 | 3571 | 3571 |
| Non-hydrogen atoms | 142230 | 140119 | 140236 |
| Waters | 11962 | 9892 | 10089 |
| Ions | 392 | 390 | 390 |
| Ligands | 3 | 3 | 3 |
| **Isotopic B factors (Å$^2$)** | | | |
| Protein | 35.0 | 40.5 | 36.9 |
| RNA | 40.1 | 44.5 | 42.7 |
| Waters | 28.7 | 29.4 | 27.4 |
| Ligands | 22.6 | 31.7 | 29.6 |
| **R.m.s. deviations** | | | |
| Bond lengths (Å) | 0.014 | 0.014 | 0.014 |
| Bond angles (°) | 1.929 | 1.867 | 1.872 |
| **Validation** | | | |
| Molprobity score | 0.85 | 0.78 | 0.78 |
| Clashscore | 1.29 | 0.91 | 0.95 |
| **Ramachandran plot (%)** | | | |
| Favored | 98.33 | 98.02 | 98.15 |
| Allowed | 1.65 | 1.94 | 1.82 |
| Outliers | 0.02 | 0.03 | 0.03 |
| **RNA geometry (%)** | | | |
| Probably wrong sugar puckers | 1.46 | 1.20 | 1.37 |
| Bad bonds | 0.04 | 0.05 | 0.05 |
| Bad angles | 0.66 | 0.53 | 0.53 |
