## Supplementary Table 2 for "The chemical landscape of the human ribosome at 1.67 Å resolution"

| rRNA | Position | Natchiar *et al.*[25] | Taoka *et al.*[21] (MS) | This study | CH...O interaction acceptor | Hydrophobic interaction | H-bond partner |
|---|---|---|---|---|---|---|---|
| 28S | 237 | xp6G | | | | | |
| 28S | 373 | Gm | | | | | |
| 28S | 398 | Am | Am | Am | Water / Protein carbonyl | | |
| 28S | 400 | | Am | Am | | LEU side chain | |
| 28S | 729 | m2G | | | | | |
| 28S | 978 | m2G | | | | | |
| 28S | 1316 | Gm | Gm | Gm | PO4 / Water | Ribose C5’ | |
| 28S | 1322 | m1A | m1A | m1A | PO4 | | |
| 28S | 1323 | | Am | | | | |
| 28S | 1326 | Am | Am | Am | Water / Ribose O4' | | |
| 28S | 1340 | | Cm | Cm | Ribose O4' | TRP / LYS side chains | |
| 28S | 1348 | xp4U | | | | | |
| 28S | 1456 | m3C | | | | | |
| 28S | 1517 | m2G | | | | | |
| 28S | 1522 | Gm | Gm | Gm | Ribose O4' | Ribose C5’ | |
| 28S | 1524 | Am | Am | Am | Ribose O4' / PO4 / Water | | |
| 28S | 1534 | Am | Am | Am | Ribose O4' | Base C6 | |
| 28S | 1536 | | Ψ | Ψ | | | Water |
| 28S | 1574 | xp6G | | | | | |
| 28S | 1582 | Ψ | Ψ | Ψ | | | Base O2 |
| 28S | 1605 | m7G | | | | | |
| 28S | 1625 | Gm | Gm | Gm | PO4 / TYR side chain | TYR side chain | |
| 28S | 1659 | xp4U | | | | | |
| 28S | 1677 | Ψ | Ψ | Ψ | | | PO4 |
| 28S | 1683 | Ψ | Ψ | Ψ | | | Water |
| 28S | 1744 | | Ψ | Ψ | | | Water |

| | | | | | | | |
|---|---|---|---|---|---|---|---|
| 28S | 1760 | | Gm | ND | | | |
| 28S | 1773 | | Um | ND | | | |
| 28S | 1779 | | Ψ | ND | | | |
| 28S | 1781 | | Ψ | Ψ | | | Water |
| 28S | 1782 | | Ψ | Ψ | | | Water |
| 28S | 1792 | | Ψ | Ψ | | | Water |
| 28S | 1797 | xe7G | | | | | |
| 28S | 1860 | m1Ψ | Ψ | Ψ | | | Water |
| 28S | 1862 | | Ψ | Ψ | | | Water |
| 28S | 1866 | m3U | | | | | |
| 28S | 1871 | Am | Am | Am | Water | | |
| 28S | 1881 | | Cm | | | | |
| 28S | 1883 | Gm | | | | | |
| 28S | 1909 | xp7G | | | | | |
| 28S | 2050 | Gm | | | | | |
| 28S | 2297 | xe7G | | | | | |
| 28S | 2351 | | Cm | Cm | | MET side chain | |
| 28S | 2363 | Am | Am | Am | Water | | |
| 28S | 2364 | Gm | Gm | Gm | Ribose O4' | | |
| 28S | 2365 | Cm | Cm | Cm | Water / Ribose O4' | Base C6 | |
| 28S | 2380 | m6G | | | | | |
| 28S | 2401 | Am | Am | Am | Water / Protein carbonyl | | |
| 28S | 2415 | | Um | Um | ND | ND | |
| 28S | 2422 | Cm | Cm | Cm | Water | | |
| 28S | 2424 | Gm | Gm | Gm | Ribose O4' | Ribose C1’ | |
| 28S | 2508 | Ψ | Ψ | Ψ | | | PO4 |
| 28S | 2522 | m7G | | | | | |
| 28S | 2754 | xp6G | | | | | |
| 28S | 2632 | | Ψ | Ψ | | | Water |

| | | | | | | | |
|---|---|---|---|---|---|---|---|
| 28S | 2773 | Gm | | | | | |
| 28S | 2786 | xp3Cm | | | | | |
| 28S | 2787 | | Am | Am | PO4 | Ribose C3’ | |
| 28S | 2804 | Cm | Cm | Cm | Ribose O4' / TYR side chain | TYR side chain | |
| 28S | 2815 | | Am | Am | PO4 | | |
| 28S | 2824 | | Cm | Cm | Water | | |
| 28S | 2837 | | Um | Um | Water / Protein carbonyl | | |
| 28S | 2835 | | Ψ | A | | | |
| 28S | 2839 | | Ψ | Ψ | | | Water |
| 28S | 2861 | Cm | Cm | Cm | Ribose O2' | Base C5 | |
| 28S | 2876 | | Gm | Gm | Water / PO4 | | |
| 28S | 3627 | | Gm | Gm | Water / PO4 | | |
| 28S | 3637 | | Ψ | Ψ | | | Water |
| 28S | 3639 | | Ψ | Ψ | | | Water |
| 28S | 3695 | | Ψ | Ψ | | | Water |
| 28S | 3701 | Cm | Cm | Cm | PO4 | | |
| 28S | 3715 | Ψ | Ψ | Ψ | | | Base O2 |
| 28S | 3718 | Am | Am | Am | Ribose O2' | | |
| 28S | 3723 | Am | Am | Am | ND | ND | |
| 28S | 3729 | Ψ | Ψ | Ψ | | | PO4 |
| 28S | 3734 | | Ψ | ND | | | |
| 28S | 3744 | | Gm | Gm | Ribose O4' | | |
| 28S | 3758 | | Ψ | ND | | | |
| 28S | 3760 | | Am | ND | | | |
| 28S | 3762 | m1Ψ | Ψ | ND | | | |
| 28S | 3764 | Ψ | Ψ | ND | | | |
| 28S | 3768 | | Ψ | ND | | | |
| 28S | 3770 | | Ψ | ND | | | |
| 28S | 3782 | m5C | m5C | m5C | Water | | |

| | | | | | | | |
|---|---|---|---|---|---|---|---|
| 28S | 3785 | Am | Am | Am | | Base C5 / CM’ | |
| 28S | 3792 | Gm | Gm | Gm | Water | | |
| 28S | 3808 | | Cm | Cm | Ribose O4' | | |
| 28S | 3818 | | Ψm | Ψm | Water | HIS side chain | PO4 |
| 28S | 3822 | | Ψ | ND | | | |
| 28S | 3825 | Am | Am | Am | Water | | |
| 28S | 3830 | | Am | Am | Water | | |
| 28S | 3841 | | Cm | Cm | Water | | |
| 28S | 3844 | | Ψ | Ψ | | | Water |
| 28S | 3851 | | Ψ | Ψ | | | Water |
| 28S | 3853 | | Ψ | Ψ | | | PO4 |
| 28S | 3867 | Am | Am | Am | Water / Ribose O4' | Base C2 | |
| 28S | 3869 | Cm | Cm | Cm | Ribose O4' | LYS side chain | |
| 28S | 3880 | xp7G | | | | | |
| 28S | 3884 | | Ψ | Ψ | | | Water |
| 28S | 3887 | Cm | Cm | Cm | Water | Base C2 | |
| 28S | 3897 | ac7G | | | | | |
| 28S | 3899 | ac7Gm | Gm | Gm | Water | | |
| 28S | 3909 | Cm | | | | | |
| 28S | 3920 | | Ψ | Ψ | | | Water |
| 28S | 3925 | | Um | Um | Protein carbonyl | PRO side chain | |
| 28S | 3944 | | Gm | ND | | | |
| 28S | 3959 | | Ψ | ND | | | |
| 28S | 4042 | | Gm | ND | | | |
| 28S | 4054 | | Cm | ND | | | |
| 28S | 4083 | m5U | | | | | |
| 28S | 4129 | m6G | | | | | |
| 28S | 4185 | m6G | | | | | |
| 28S | 4194 | xp4U | | | | | |

| | | | | | | | |
|---|---|---|---|---|---|---|---|
| 28S | 4196 | Gm | Gm | Gm | Ribose O2' / Ribose O4' / Water | Ribose C1’ | |
| 28S | 4220 | m6A | m6A | m6A | Ribose O2' | | |
| 28S | 4227 | | Um | Um | Ribose O4' / Water | | |
| 28S | 4228 | | Gm | Gm | Water / THR side chain | | |
| 28S | 4293 | Ψ | Ψ | Ψ | | | Ribose O2’ |
| 28S | 4296 | m1Ψ | Ψ | Ψ | | | Water |
| 28S | 4299 | | Ψ | Ψ | | | Water |
| 28S | 4306 | Um | Um | Um | Water / TYR side chain | | |
| 28S | 4312 | | Ψ | Ψ | | | Water |
| 28S | 4335 | m5C | | | | | |
| 28S | 4353 | | Ψ | Ψ | | | Water |
| 28S | 4355 | xe6G | | | | | |
| 28S | 4361 | | Ψ | Ψ | | | Water |
| 28S | 4370 | Gm | Gm | Gm | ND | ND | |
| 28S | 4371 | m2xp7G | | | | | |
| 28S | 4392 | | Gm | Gm | | Base / CM5 | |
| 28S | 4403 | Ψ | Ψ | Ψ | | | Water |
| 28S | 4415 | m1A | | | | | |
| 28S | 4420 | | Ψ | Ψ | | | PO4 |
| 28S | 4423 | | Ψ | Ψ | | | Water |
| 28S | 4431 | | Ψ | Ψ | | | Water |
| 28S | 4442 | Ψ | Ψ | Ψ | | | Water |
| 28S | 4447 | m5C | m5C | m5C | Ribose O4' | CM’ | |
| 28S | 4450 | Ψ | | | | | |
| 28S | 4456 | | Cm | Cm | | PRO side chain | |
| 28S | 4457 | | Ψ | Ψ | | | Water |
| 28S | 4471 | | Ψ | Ψ | | | Water |
| 28S | 4472 | m6G | | | | | |
| 28S | 4483 | m4C | | | | | |

| | | | | | | | |
|---|---|---|---|---|---|---|---|
| 28S | 4493 | | Ψ | Ψ | | | Ribose O2’ |
| 28S | 4494 | Gm | Gm | Gm | Ribose O4' | VAL side chain | |
| 28S | 4498 | | Um | Um | Ribose O4' | Base C8 | |
| 28S | 4499 | | Gm | Gm | Water / Ribose O2' | | |
| 28S | 4500 | Ψ | Ψ | Ψ | | | Base O4 |
| 28S | 4521 | | Ψ | Ψ | | | Water |
| 28S | 4523 | Am | Am | Am | PO4 | | |
| 28S | 4529 | m6G | | | | | |
| 28S | 4530 | m3U | m3U | m3U | Water | | |
| 28S | 4531 | Ψ | | | | | |
| 28S | 4532 | | Ψ | Ψ | | | Water |
| 28S | 4536 | Cm | Cm | Cm | Ribose O4' | CM’ | |
| 28S | 4550 | m7G | | | | | |
| 28S | 4552 | | Ψ | Ψ | | | Water |
| 28S | 4564 | m7A | | | | | |
| 28S | 4571 | Am | Am | Am | Water | Ribose C1’ | |
| 28S | 4576 | | Ψ | Ψ | | | Water |
| 28S | 4579 | | Ψ | Ψ | | | Water |
| 28S | 4590 | | Am | Am | Water | | |
| 28S | 4597 | m3U | | | | | |
| 28S | 4618 | | Gm | Gm | Water | | |
| 28S | 4620 | Um | Um | Um | Water | Ribose C5’ / PRO side chain | |
| 28S | 4623 | Gm | Gm | Gm | Ribose O4' | | |
| 28S | 4628 | Ψ | Ψ | Ψ | | | Water |
| 28S | 4636 | Ψ | Ψ | ND | | | |
| 28S | 4637 | Gm | Gm | Gm | Base O2 | | |
| 28S | 4671 | m4C | | | | | |
| 28S | 4673 | | Ψ | Ψ | | | Water |
| 28S | 4689 | | Ψ | Ψ | | | Water |

| | | | | | | |
|---|---|---|---|---|---|---|
| 28S | 4690 | ac7G | | | | |
| 28S | 4870 | Gm | | | | |
| 28S | 4872 | m2G | | | | |
| 28S | 4972 | | Ψ | Ψ | | Water |
| 28S | 5001 | | Ψ | Ψ | | Water |
| 28S | 5010 | | Ψ | Ψ | | Water |
| 5.8S | 14 | Um | Um | | | |
| 5.8S | 55 | | Ψ | Ψ | | Water |
| 5.8S | 69 | | Ψ | Ψ | | Water |
| 5.8S | 75 | | Gm | Gm | MET side chain | |
| Total | | 102 | 137 | 117 | | |
