## Supplementary Figures for "The chemical landscape of the human ribosome at 1.67 Å resolution"

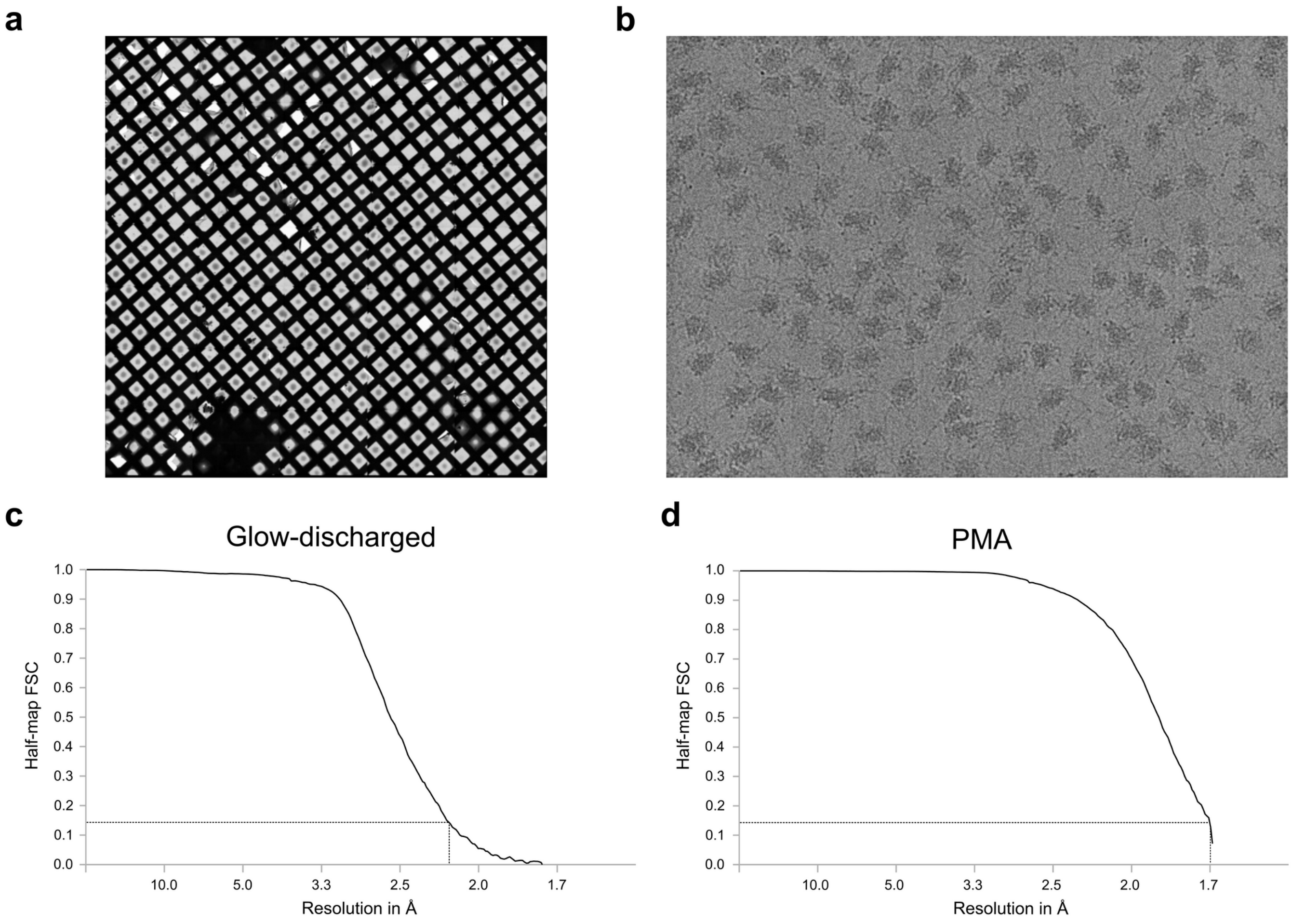


**Supplementary Figure 1: FSC curves for datasets from glow-discharged and PMA-treated grids.** **a.** Atlas view of PMA-treated grid showing homogeneous thin ice across the grid. **b.** Representative motion-corrected micrograph acquired at -1.5 µm defocus from the PMA-treated grid dataset. **c.** Gold standard FSC curve for the glow-discharged dataset. **d.** Gold standard FSC curve for the PMA dataset.

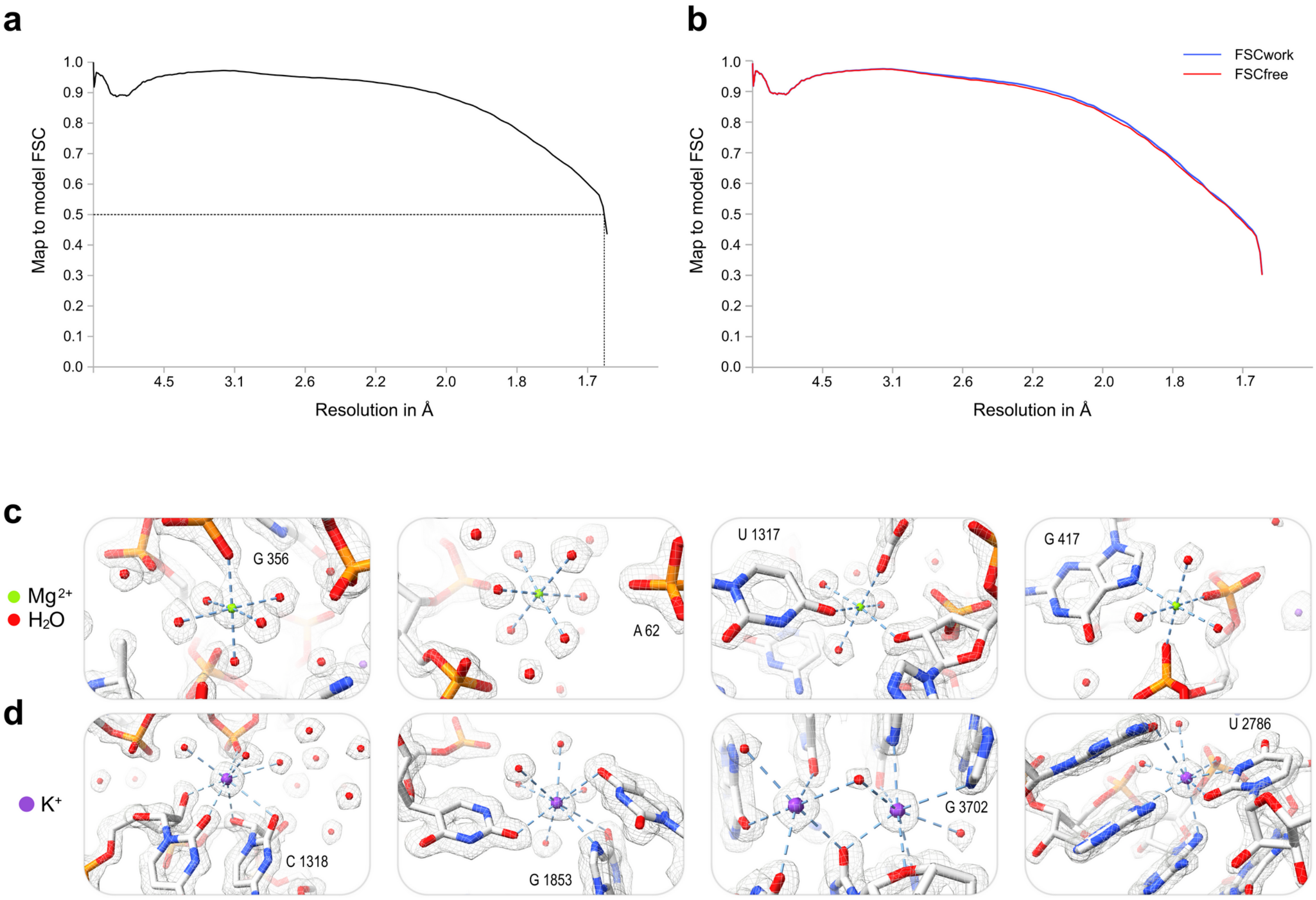


**Supplementary Figure 2: Human 60S structure refinement and validation from the PMA dataset.**

**a.** Map to model FSC curve (threshold = 0.5). **b.** Cross-validation against overfitting. **c-d.** Examples of $Mg^{2+}$ (**c**) and $K^{+}$ (**d**) ion coordination (light blue dashed lines).

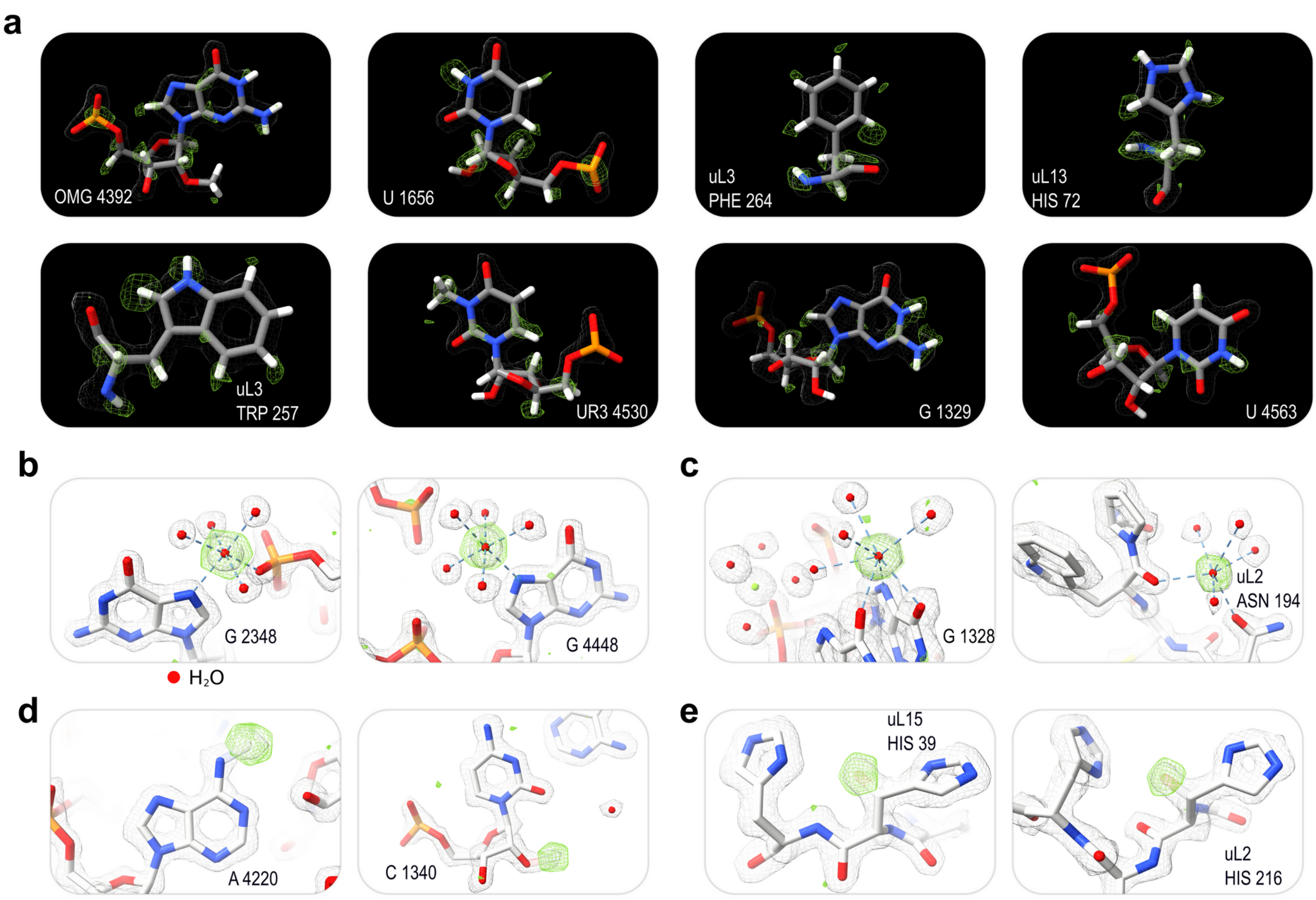


**Supplementary Figure 3: Direct observation of hydrogen atoms and rRNA modifications**

**a.** Examples of signal for hydrogen atoms in rRNA and amino acid residues observed in Fo-Fc difference map**. b-e.** Fo-Fc difference maps reveal inaccurate structural modelling. Positive difference signal (green mesh) highlights $Mg^{2+}$ (**b**) and $K^{+}$ (**c**) ions incorrectly modelled as waters, unmodelled rRNA methylation (**d**) or ribosomal protein histidine beta-hydroxylation (**e**). Hydrogen atoms are depicted as white sticks for visualisation purposes only and were not present in the structure used to calculate the Fo-Fc map. Unmodeled modifications are shown as semi-transparent sticks.

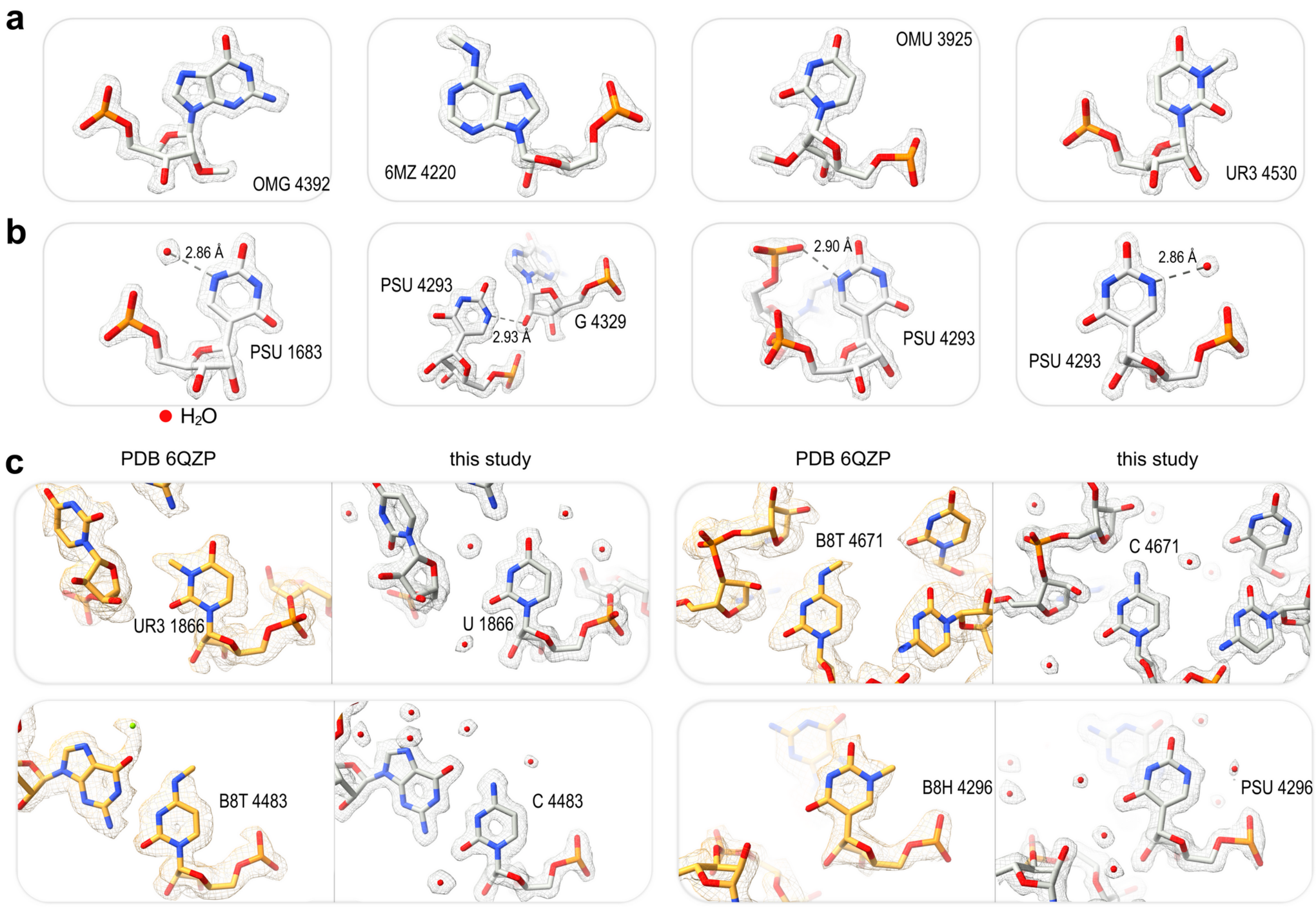


**Supplementary Figure 4: Directly visualising rRNA modifications.**

**a-b.** Examples of rRNA methylation (**a**) and pseudouridylation (**b**). Hydrogen bonds (grey dashed lines) and distances between the N1 atom of pseudouridine and neighbouring heteroatoms are indicated. **c.** Density for current model (right) compared with that used to support previously proposed human rRNA modifications (PDB 6QZP; EMD-4245) (left, orange). OMG, 2'-O-methylguanosine. 6MZ, N6-methyladenosine. OMU, 2'-O-methyluridine. UR3, 3-methyluridine. B8T, 4-methylcytosine. B8H, 1-methyluridine.

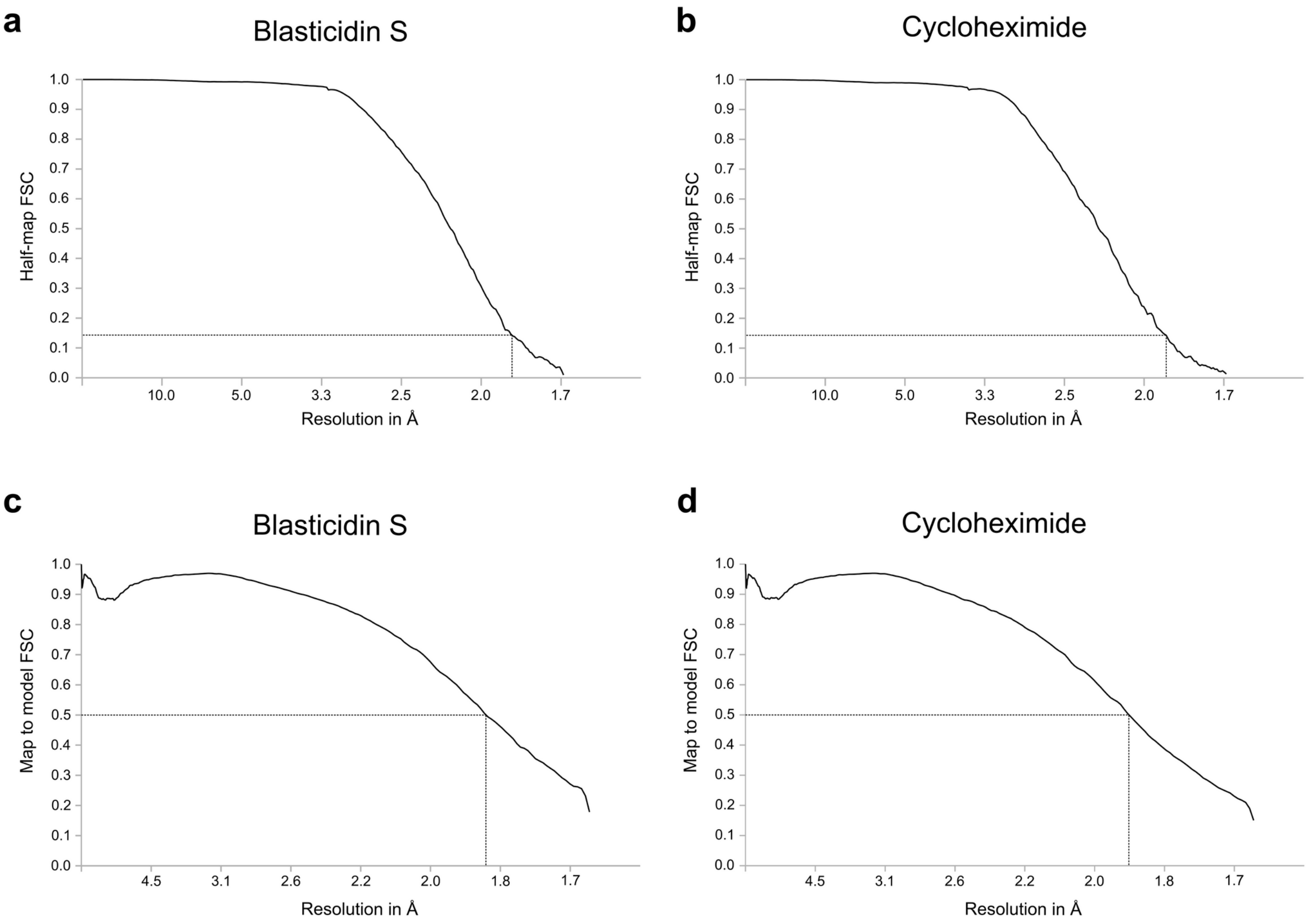


**Supplementary Figure 5: FSC curves for human 60S-inhibitor complexes. a-b.** Gold standard FSC curves for datasets corresponding to human 60S subunits bound to blasticidin S (**a**) and cycloheximide (**b**). **c-d.** Map to model FSC curves for datasets corresponding to human 60S subunits bound to blasticidin S (**c**) and cycloheximide (**d**) (threshold = 0.5).

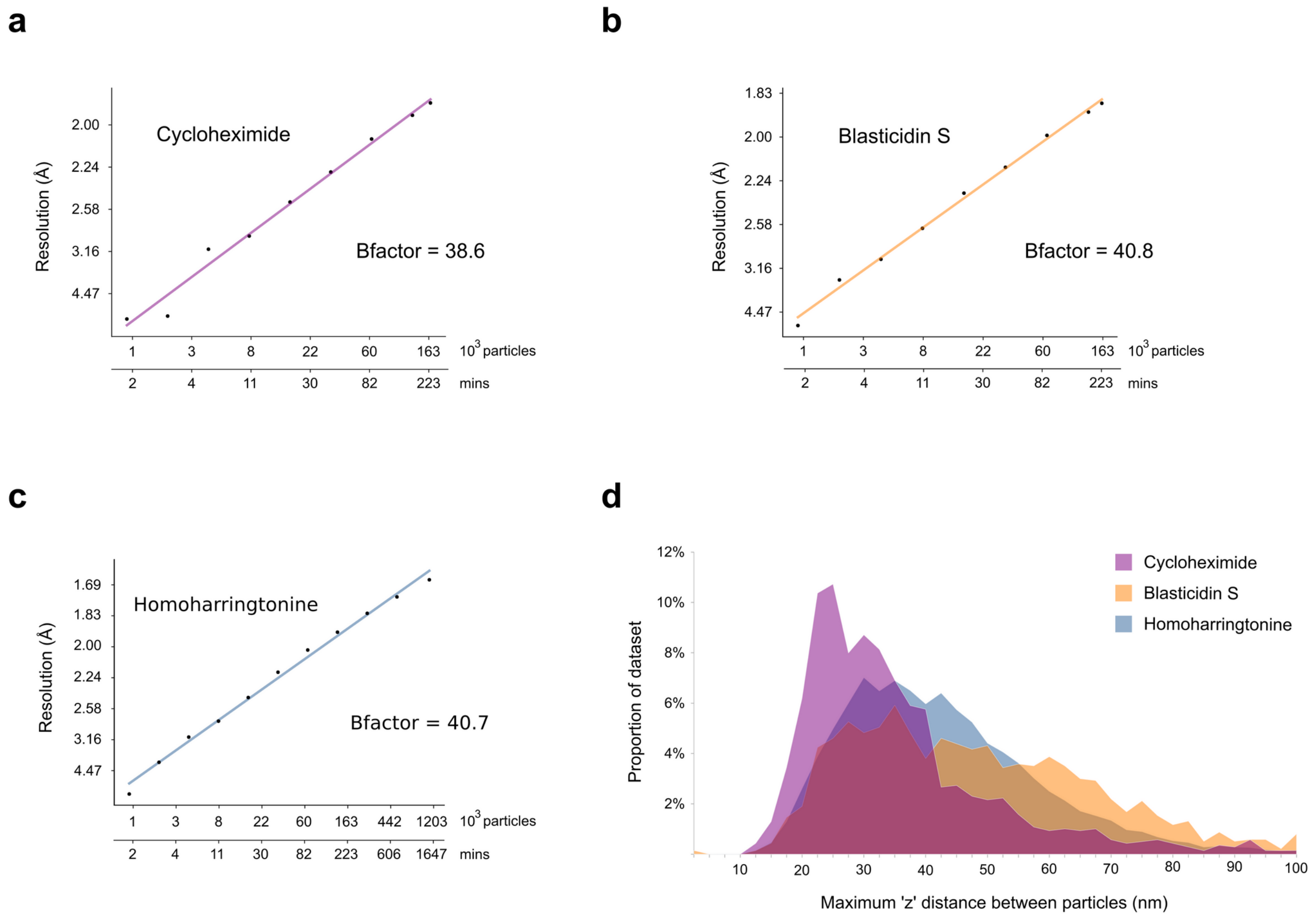


**Supplementary Figure 6: B-factor plots and distribution of micrographs by maximum inter-particle 'z' distance. a-c.** Rosenthal-Henderson 'B-factor' plots for cycloheximide (**a**), blasticidin S (**b**), and homoharringtonine (**c**) complexes. **d.** Micrograph distribution as a function of the range of refined defocus values for the particles.

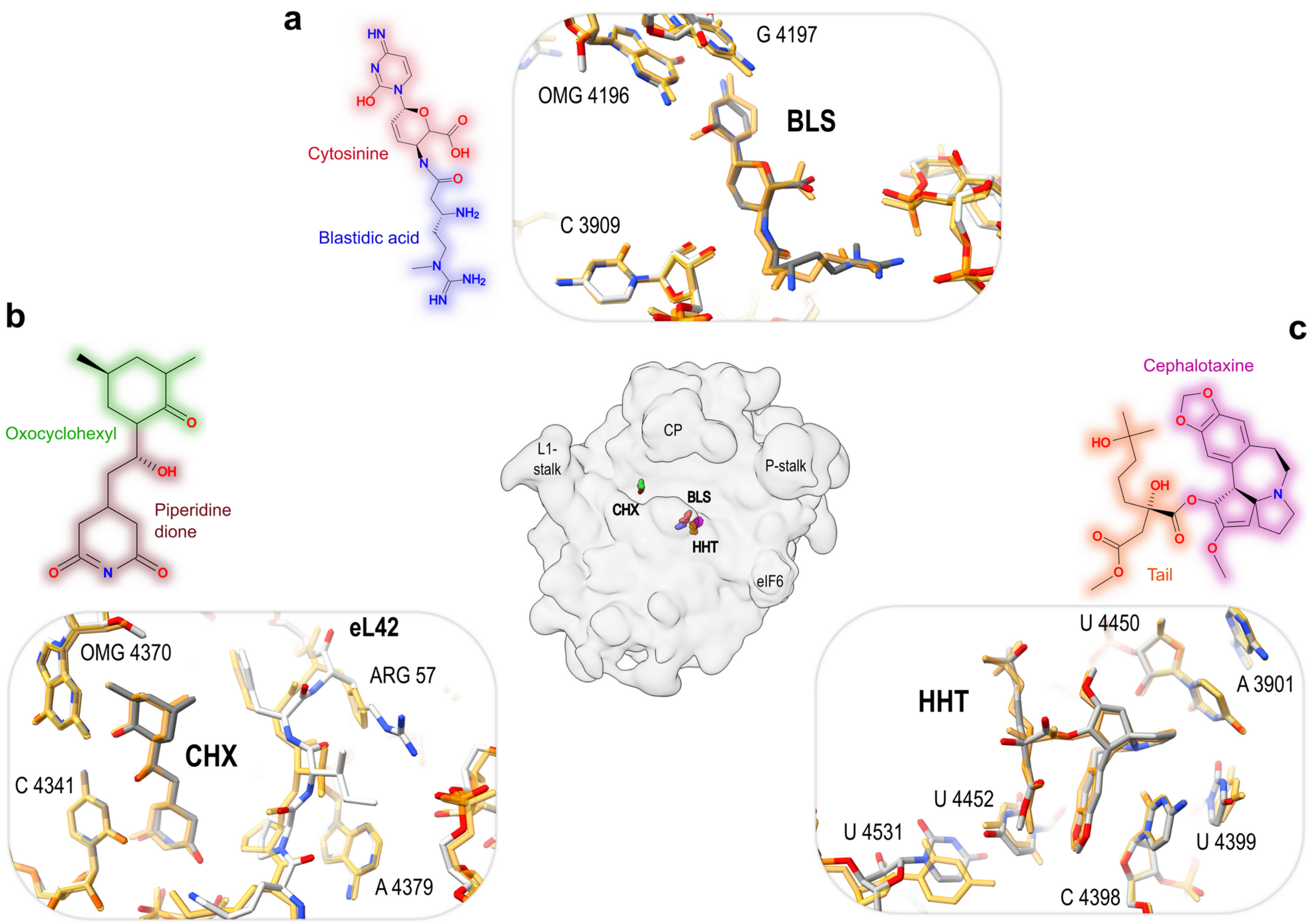


**Supplementary Figure 7: 60S inhibitor structures and interactions.**

**Central panel.** Localisation of inhibitors bound to the 60S ribosomal subunit. **a-c.** 2D chemical structures coloured by chemical group and superposition of corresponding yeast 80S-ligand complexes on the current structures for blasticidin S (PBD ID 4U56) (**a**), cycloheximide (PDB ID 4U3U) (**b**), and homoharringtonine (PDB ID 4U4Q) (**c**). BLS, blasticidin S. CHX, cycloheximide. HHT, homoharringtonine.

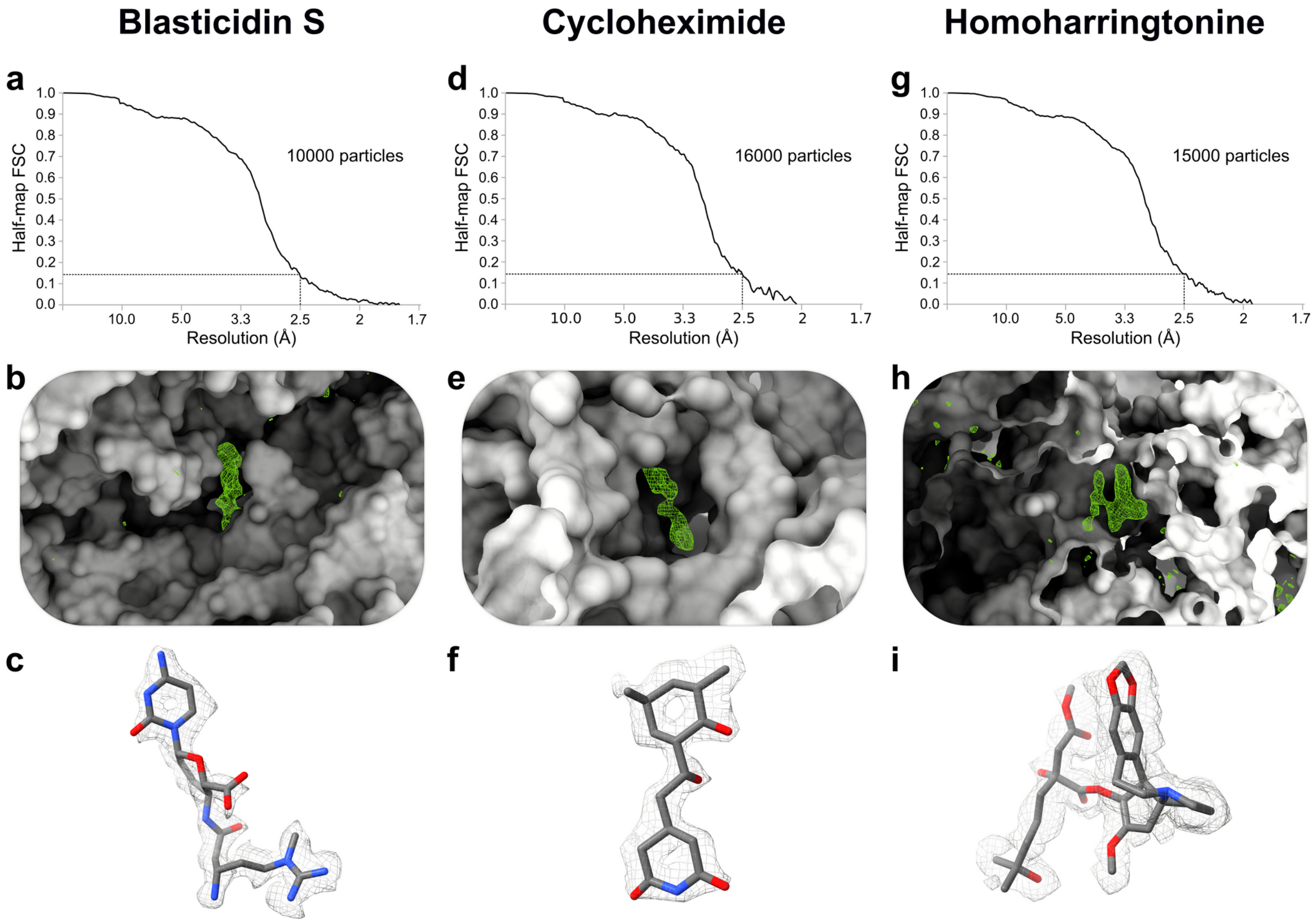


**Supplementary Figure 8: Modelling of bound 60S inhibitors in cryo-EM maps reconstructed from small datasets.**

**a-j.** Gold standard FSC curves, Fo-Fc difference maps comparing inhibitor-bound versus apo 60S reconstructions with positive signal difference shown as green mesh, 2D chemical structure of inhibitors and fitting into 2.5 Å resolution map for blasticidin S (**a-d**), cycloheximide (**e-h**) and homoharringtonine (**i-l**) respectively**.**
